# Endogenous fine-mapping of functional regulatory elements in complex genetic loci

**DOI:** 10.1101/2023.05.06.539696

**Authors:** Ke Zhao, Yao Zhou, Chengyue Wu, Jianhua Wang, Hongcheng Yao, Xin Cheng, Lin Zhao, Wei Wang, Xinlei Chu, Xianfu Yi, Yupeng Chen, Miaoxin Li, Wange Lu, Kexin Chen, Pak Chung Sham, Mulin Jun Li

## Abstract

The vast majority of genetic loci associated with polygenic complex traits are located in non-coding regions of the human genome. However, many of these regions exhibit high-order gene regulatory relationships and complicated linkage disequilibrium (LD) configurations, which bring challenges to accurately identify causal variants and their target genes controlling specific molecular processes or traits. We employed multiplexed single-cell CRISPR interference and activation perturbations to explore the links between *cis*-regulatory element (CRE) and target gene expression within tight LD in the endogenous chromatin context. We validated the prevalence of multiple causality in perfect LD (pLD) for independent expression quantitative trait locus (eQTL), and revealed fine-grained genetic effects on gene expression within pLD. These effects are difficult to decipher using conventional eQTL fine-mapping or to predict via existing computational methods. We found that nearly half of the casual CREs lack classical epigenetic markers, potentially affecting gene expression through hidden regulatory mechanisms. Integrative analysis on different types of perturbation effects suggested a high regulatory plasticity of the human genome. These findings will propel further in-depth exploration of functional genomic elements, facilitating a more comprehensive understanding of gene expression regulatory patterns and the development of complex traits.

## Introduction

Identifying fine-grained regulatory elements and complex trait/disease causal regulatory variants on the human genome is a significant challenge in the current fields of functional genomics and genetics. Years of functional genomic profiling and expression quantitative trait locus (eQTL) studies have identified numerous *cis*-regulatory elements (CREs) and expression-associated alleles in nearly a hundred human tissues/cells (*1–3*). However, accurately pinpointing which CREs or even single allele(s) can modulate gene expression under specific biological conditions remains difficult. This is especially true for complex genetic loci, where complicated CRE-target gene relationships and linkage disequilibrium (LD) contamination make it computationally and experimentally challenging to precisely locate all causal elements and their target genes (*4*). Furthermore, genome-wide association studies (GWASs) have shown that the majority of genetic loci associated with complex traits and diseases are located in non-coding regions of the genome; however, colocalization analysis with eQTLs revealed only a limited (8%-25%) proportion of shared genetic loci (*5–7*). This has sparked debate within the field over whether a substantial portion of trait/disease-causal variants may not cause phenotypic development by affecting gene expression.

A variety of large-scale, high-throughput experimental methods have been used to systematically evaluate the regulatory potential of human genomic sequences and allele-specific effects. Firstly, exogenous and episomal massively parallel reporter assays (MPRAs) are highly efficient in identifying functional regulatory sites and alleles in various cells (*8, 9*). Consistent with computational simulations (*4*), a recent study employed MPRA to deeply assess high LD variants within independent eQTL signals, systematically validating the widespread presence of multiple allelic effects in tight LD (*10*). However, these experiments struggle to evaluate genetic effects and relationships with specific molecular phenotypes (e.g., target gene expression) in the endogenous chromatin environment of local genetic loci. Additionally, using CRISPR base editing technology, researchers have been able to study the links between variants and complex phenotypes in targeted genomic regions (*11–14*). However, due to the restrictions of editing preference and efficiency in mammalian cells, most of these strategies are not suitable for fine-mapping causal genetic loci affecting molecular phenotypes. Finally, some studies employ CRISPR-based chromatin perturbations to exhaustively characterize regulatory sites and their causal relationships with particular gene expression(s) through tiling screening (*15–18*), but these studies have focused solely on a small number of target genes.

Multiplexed single-cell CRISPR perturbations provide technical support for systematically studying the regulatory relationships between genetic loci and fine-grained molecular phenotypes in an endogenous cellular environment (*19–21*). However, these studies heavily rely on prior knowledge (e.g., specific epigenetic markers) for the selection of genetic loci and employ solely a single type of CRISPR perturbation. Whether such technologies can accurately and unbiasedly pinpoint causal CREs that regulate gene expression within complex genetic loci (e.g., ultra-high LD region) is a scientific question worth exploring. In this study, we leveraged multiplexed single-cell perturbations, through both CRISPR interference and activation, to validate the ubiquity of multiple causal CREs in perfect LD (pLD) within independent eQTL signal and to interrogate their regulatory relationships with target genes. We have demonstrated that endogenous perturbations can reveal intricate genetic effects and gene expression regulatory patterns, which are challenging to identify through conventional eQTL mapping approaches. Moreover, we found that current computational methods struggle to accurately predict the effects of many endogenous perturbations on gene expression, and further investigation suggested that nearly half of the casual CREs lack classical epigenetic markers, potentially influencing gene expression through unique regulatory mechanisms. Lastly, by comparing various types of CRISPRi/a perturbations, we have shed light on the regulatory plasticity of the human genome through a distinct perspective.

## Results

### Comprehensive discrimination of independent eQTL effect at perfect LD (pLD) using multiplexed single-cell CRISPR perturbations

Given fine-mapping causal allele(s) among perfectly correlated variants is either computationally intractable or experimentally costly, we designed a series of extreme scenarios and applied a single-cell perturbation strategy to provide unbiased interrogation of expression-modulating causal effects at native genomic settings. Generally, we first leveraged statistical fine-mapping on GTEx V8 whole-genome sequencing (WGS)-based eQTLs (670 whole blood samples) to nominate independent signals. Additionally, we collected and fine-mapped six WGS-based blood-derived eQTL datasets from Geuvadis (445 lymphoblastoid cell samples), BLUEPRINT (190 monocyte samples, 196 neutrophil samples and 165 CD4+ T cell samples), and TwinsUK (523 lymphoblastoid cell samples and 246 whole blood samples) (Table S1), and required that independent signal is reproducible in at least one additional dataset. Ultimately, we integrated all fine-mapped variants in seven eQTL datasets to retain independent signals which contain at least two undistinguishable lead eQTL variants (LEVs, with equal causal probability) in pLD (Fig. 1A and Fig. S1A, see details in Methods). As large-scale base editing screens, such as prime editing (*13*), coupled with transcriptome readouts are currently impractical, we performed multiplexed regional perturbations at the single-cell level to investigate the causal relationships and multiplicity underlying eQTL effects within these complex signals (Fig. 1A, see details in Methods).

**Fig. 1.**
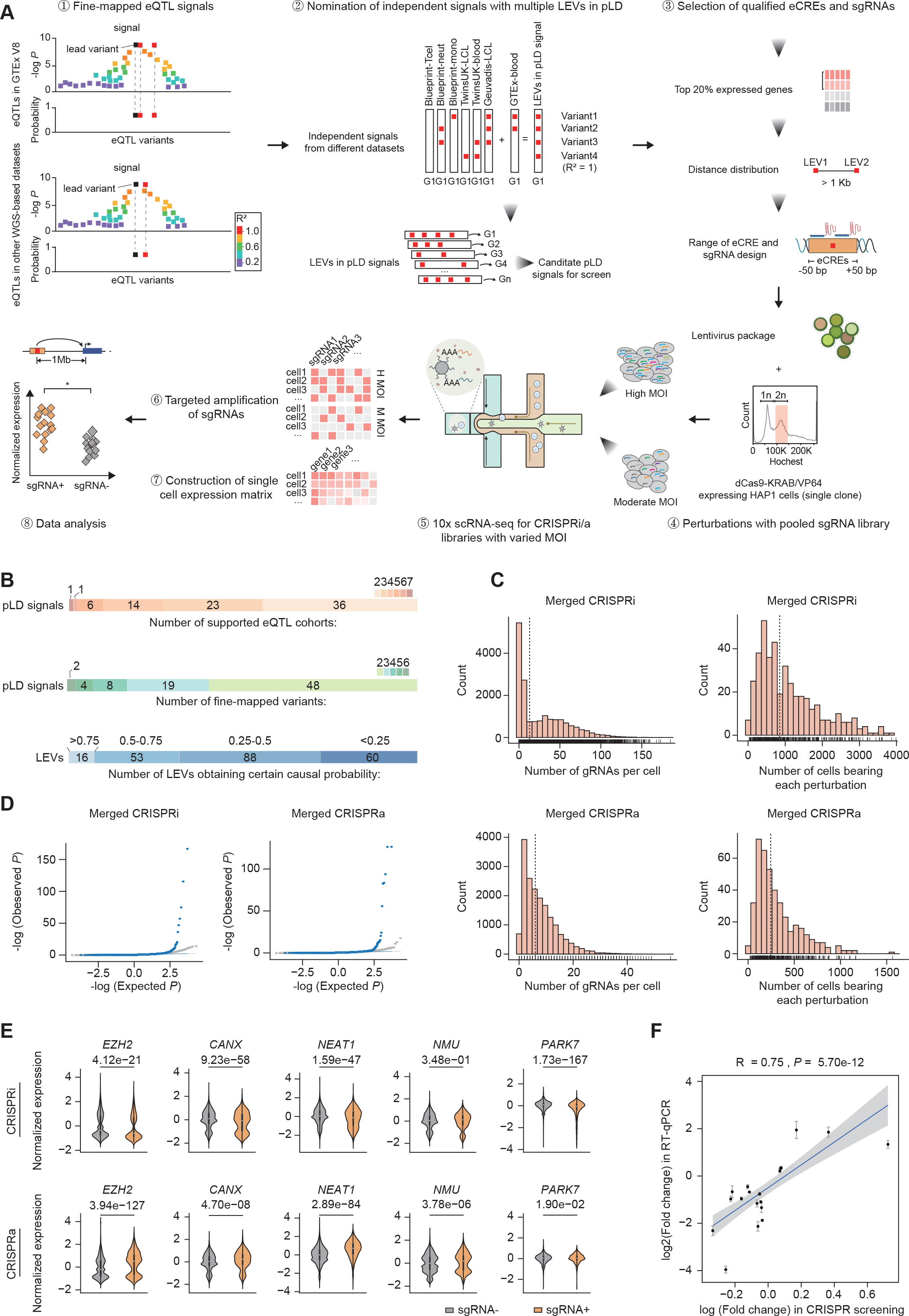
**Identification of causal eQTL-tagged CREs in pLD regions using multiplexed single-cell CRISPRi/a perturbations. a**, Schematic of the multiplexed single-cell CRISPRi/a perturbations used to screen for causal eCREs tagged by eQTL variants in pLD signals and to systematically study the regulatory relationships between genetic loci and gene expression levels. LEV: lead eQTL variants with equal fine-mapped causal probability; eCRE: LEV-tagged CRE (± 50 bp region of LEVs); pLD: perfect LD (R^2^=1); MOI: multiplicity of infection; Blueprint-Tcel, Blueprint-neut and Blueprint-mono: fine-mapped eQTL data of CD4+ T cell samples, neutrophil samples and monocyte samples from BLUEPRINT project; TwinsUK-LCL and TwinsUK-blood: fine-mapped eQTL data of lymphoblastoid cell samples and whole blood samples from UK10K TwinsUK project; Geuvadis-LCL: fine-mapped eQTL data of lymphoblastoid cell samples from Geuvadis project; GTEx V8-blood: fine-mapped eQTL data of whole blood samples from GTEx V8 project. **b**, Summary information for the selected qualified LEVs and their corresponding pLD signals, including the number of independent eQTL datasets that support corresponding pLD signals, the number of fine-mapped LEVs within corresponding pLD signals, and the causal probability (mean score of different datasets, estimated via CaVEMaN) distribution of selected fine-mapped LEVs. **c**, The number of gRNAs per cell and the number of cells bearing each perturbation in CRISPRa/i screen, based on integration of data from both high or moderate MOIs. **d**, Quantile-quantile plot comparing observed versus expected P of eCRE-targeting sgRNAs (blue) and non- targeting control (NTC) sgRNAs (gray; down-sampled) associated with gene expression. **e**, Results of positive control sgRNAs in the CRISPRi and CRISPRa perturbations. sgRNAs targeting *EZH2*, *CANX*, *NEAT1*, *PARK7* are positive control targeting promoter of genes, and sgRNA effect on *NMU* is associated with an enhancer. **f**, Consistence analysis of perturbation results and RT-qPCR results for the 20 randomly selected causal eCREs-perturbGene pairs.

To achieve effective discrimination of single-cell expression perturbations via CRISPR, we focused on eQTL genes (eGenes) which are highly (top 20%) expressed in a human near-haploid leukemia cell line (HAP1), and excluded unqualified LEVs (e.g., proximal variants or protein-coding/splicing-altering variants) in each signal (Fig. S1A and Fig.S1B, see details in Methods). Thus, 81 independent eQTL signals (reproducible in 2 to 7 additional datasets) were finally selected, and each contains more than a single qualified LEV within pLD (range of 2 to 6). Finally, the perturbation library incorporates 217 LEVs (Fig. 1B and Table S2). Two or three single guide RNAs (sgRNAs) were designed to target each LEV-tagged region, and additional two sets of control sgRNAs were also used, including 39 non-targeting control (NTC) sgRNAs as negative control and 11 previously validated sgRNAs as positive control (*21*). The sequence of 472 designed sgRNAs was cloned into a lentiviral CROP-seq-opti vector (Table S3) (*22*). The quality of the constructed sgRNA plasmid library was validated and was found to be of high quality, as evidenced by the coverage rate of greater than 99% (Fig. S1C) and a low degree of uniformity, with the top 90^th^ to 10^th^ sgRNA having a representation difference of less than 10-fold (Fig. S1D). Consistency in sgRNA distribution within plasmid library was also observed upon viral transduction at varied multiplicity of infection (MOI) (Fig. S1E).

To maximize the detection power of CRISPR-based perturbations for fine-mapped LEV- tagged CRE (eCRE, ± 50 bp region of LEVs) discovery, we used both CRISPR interference (CRISPRi, a nuclease-deactivated Cas9 tethered to the KRAB repressor domain, dCas9-KRAB) and CRISPR activation (CRISPRa, a dCas9 tethered to the transcriptional activator VP64, dCas9-VP64) systems (Fig. S1F). Previous studies have demonstrated that the introduction of multiple perturbations per cell can substantially augment the statistical power to identify causal relationships between CREs and their target genes in a cost-efficient manner (*21, 23, 24*). Thus, lentiviral vectors containing our sgRNA library were then transduced into selected monoclonal cell lines at either moderate or high MOI (Fig. S1F-S1I). This approach allowed us to comprehensively investigate the causal eCREs and their target genes among multiple eQTL signals, while minimizing genetic heterogeneity in our experimental system. After a 14-day cell culture for effective CRISPRi/a perturbation, the transcriptomes of 14,481/15,296 (CRISPRi) and 17,303/17,709 (CRISPRa) single cells were profiled with four 10x single-cell RNA sequencing (scRNA-seq) libraries. Targeted amplification of sgRNAs from cDNA in these perturbation libraries (*25*) suggested that the number of sgRNAs per cell and the number of cells per perturbation was increased as MOI increases in both CRISPRi and CRISPRa perturbations (Fig. S1J and S1K). Joint analysis of data under different MOIs revealed a median of 13 sgRNAs per cell and a median of 850 cells bearing each perturbation with CRISPRi, and a median of 6 sgRNAs per cell and a median of 245 cells bearing each perturbation with CRISPRa, respectively (Fig. 1C).

We applied a unified normalization-association framework, Normalisr (*26*), to analyze the relationship between each eCRE and nearby expressed genes (± 1 Mb) (see details in Methods). Quantile-quantile plots indicated an excess of significant associations of sgRNAs targeting eCRE compared with NTCs in all conditions (132 and 34 significant pairs of eCRE and its perturbed gene (perturbGene) in merged CRISPRi (Table S4) and CRISPRa (Table S5) screenings respectively, FDR < 0.2). The perturbation with high MOI achieved higher power than moderate MOI (Fig. 1D), and the perturbation effects from significant hits between moderate and high MOIs showed good agreement (Fig. S2A and Fig. S2B). Besides, we observed significant consistency between the results of the perturbation experiments under different MOI conditions (R = 0.23, *P* = 0.0034) (Fig. S2C). To further explore the influence factors determining the statistical power of multiplexed single-cell CRISPR perturbations, we simulated several scRNA-seq datasets with various perturbation conditions, including effective sgRNAs per cell and total captured cells in each perturbation (see details in Methods). The results demonstrated that, consistent with the real experiments, increasing the number of perturbations in each cell led to better statistical power. However, simply increasing the number of captured cells in scRNA-seq only slightly promoted statistical power (Fig. S2D). These findings suggest that, for perturbation effects that are sparse in terms of their downstream consequences, multiplexed sgRNAs will offer powerful and cost-efficient solution in single-cell CRISPR perturbations.

To validate the screening results, we first confirmed the effects of all positive control sgRNAs, which were highly concordant with the literature report (Fig. 1E). Additionally, we randomly selected 20 groups of sgRNAs targeting causal eCRE and performed individual CRISPRi or CRISPRa perturbations, followed by reverse-transcription qPCR (RT-qPCR), to confirm the effect of sgRNAs on their pertubGenes, with no-targeting sgRNAs as control. As expected, a positive correlation (R = 0.75, *P* = 5.70e-12, T-test) between the RT-qPCR results and the effect size observed in multiplexed single-cell CRISPR perturbations was found (Fig. 1F and Fig. S3). This results further demonstrate the credibility of our single-cell perturbation screens in testing the effects of eCREs on their potential target genes.

### Multiple causal *cis*-effects and target gene configurations underlie complex genetic associations on gene expression

Based on the significant hits in both CRISPRi/CRISPRa screens, we first inquired whether the presence of multisite *cis*-regulation and multiplicity of target genes in pLD is prevalent. Our results showed that, over 70% of the investigated pLD signals contained at least one significant causal eCRE. Among these significant pLD signals, 49.1% of them showed multiple (2 to 5) causal eCRE (44.2% in CRISPRi and 11% in CRISPRa, respectively), suggesting a high proportion of multisite *cis*-regulation. Besides, over half (52.6%) of these pLD signals had multiple target genes, with the maximum being greater than four, while standalone CRISPRi and CRISPRa analyses revealed proportions of 51.9% and 14.8%, respectively. Additionally, we found that the perturbGenes from 39% pLD signals were the closest to the associated causal eCRE, 53% were located distally, and the remaining 9% could link to both closest and distal target genes (Fig. 2A and Fig. S4A). These results emphasize the pervasive existence of multisite *cis*-regulation affecting various target genes in pLD, which complicates the identification of true causal variants in both eQTL and GWAS fine-mapping.

**Fig. 2.**
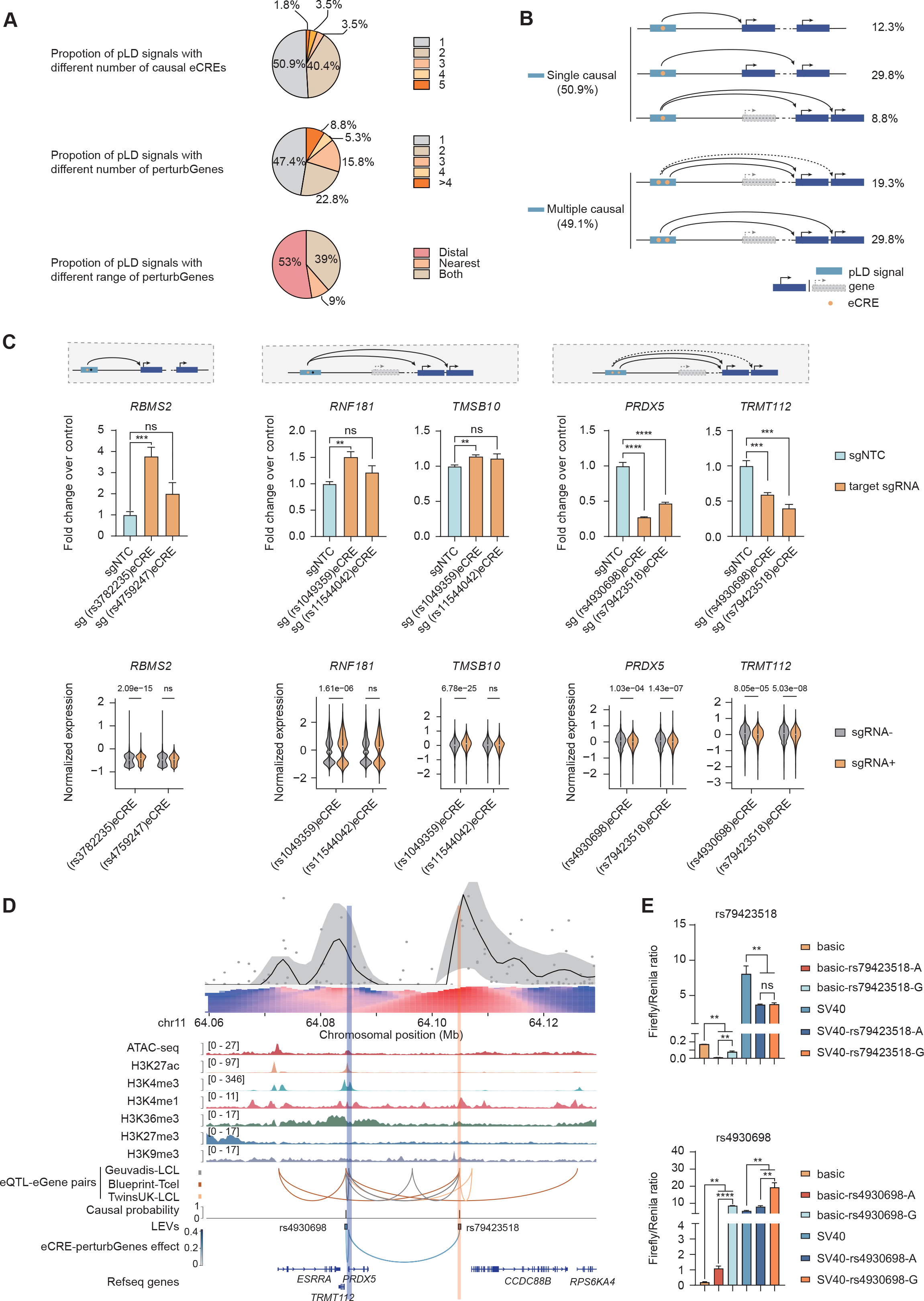
**CRISPR perturbations identify diverse regulatory patterns between causal eCREs and their target genes in pLD signals. a**, Distribution of the pLD signals containing different numbers of causal eCREs (top), regulating different numbers of perturbGenes (middle), and showing varied distances between eCREs and their perturbGenes (bottom). **b**, Patterns of causal eCREs targeting their perturbGenes in corresponding pLD signals. Genes are shown in dark blue or gray (possible nearest gene). The direction and position of TSS are indicated by arrows. pLD signals are labeled in light blue. Yellow dots represent eCREs. Arcs indicate regulatory relationships between causal eCREs and perturbGenes, while dashed lines indicate possible regulatory relationships. **c**, Validations of several representative eCRE-perturbGene pairs. Corresponding eCRE-perturbGene regulatory patterns (top), RT-qPCR analysis of perturbGene expression upon perturbation with sgRNAs targeting eCREs (middle) and differential perturbGene expressions in multiplexed CRISPR perturbations (bottom). **d**, 4C assay and epigenomic evidence indicating chromatin interactions between rs79423518-located DNA fragment (yellow shaded region) and promoter of *PRDX5* and *TMRT112* (blue shaded region). **e**, Luciferase assay results of rs4930698/rs79423518- tagged CREs and allele-specific effects of them. Error bars represent standard deviation of the mean. *P* are indicated by asterisks, with \**P* < 0.05, \*\**P* < 0.01, \*\*\**P* < 0.001, \*\*\*\**P* < 0.0001, and ns indicating not significant.

To investigate the target gene configurations underlying the complex genetic associations on gene expression, we partitioned the pLD signals containing significant eCRE- perturbGene pairs into two regulatory patterns (single causal and multiple causal) (Fig. 2B, Fig. S4B). For pLD signals with unique causal eCRE, some only associated with the nearest gene (12.3% of the total significant pLD signals) (Fig. 2B). For example, two highly linked LEVs, rs3782235 (GRCh37: chr12:56915547-G-A) and rs4759247 (12:56918834-T-G) located more than 1 Kb apart from each other. Our CIRSPRa perturbation screens revealed that only rs3782235-tagged CRE significantly affected the expression of the adjacent gene *RBMS2*, which was confirmed by RT-qPCR. Specifically, the sgRNA targeting rs3782235 significantly up-regulated the expression of *RBMS2* with a greater effect size than sgRNAs targeting rs475924, consistent with the screening results (Fig. 2C). Interestingly, a recent GWAS of blood traits (*27*) found rs3782235 was significantly associated with hematocrit percentage, suggesting that rs3782235 could be a causal variant modulating *RBMS2* expression in haematocrit-related traits. Besides, instead of the nearest gene, 29.8% significant pLD signals connected to a single distal gene via corresponding causal eCRE (Fig. 2B). Moreover, casual eCREs in a small fraction of pLD signals (8.8%) can regulate multiple target genes (Fig. 2B). For instance, CRISPRa perturbations revealed that rs1049359-tagged CRE, rather than other highly linked variant-marked eCRE in pLD, affected the expression of two distal target genes, *RNF181* and *TMSB10*, which were independently verified by RT-qPCR (Fig. 2C).

As for multiple causal patterns, 19.3% of the total significant pLD signals incorporated more than one causal eCREs targeting the same target gene(s), indicating the common phenomenon of multisite constraints on eQTL fine-mapping (*4, 10*). For example, we identified two significant eCREs (tagged by rs4930698 (chr11:64085063-G-C) and rs79423518 (chr11:64105454-G-A)) in a pLD signal that were both associated with two target genes *PRDX5* and *TRMT112*. Consistent with the CRISPRi screening results, the effect of two eCREs was confirmed through RT-qPCR, which showed that the sgRNAs targeting the corresponding eCRE significantly decreased the expression levels of both *PRDX5* and *TRMT112* (Fig. 2C). The rs4930698-tagged CRE is located in the upstream promoter region of *PRDX5* and downstream promoter region of *TRMT112*, and overlays several active chromatin marks including H3K4me3, H3K27ac and open chromatin, indicating its high regulatory potential. Besides, rs79423518-tagged CRE lies at intergenic region ∼20 Kb downstream of *PRDX5*, and obtains weak enhancer marks such as H3K4me1 and H3K27ac (Fig. 2D). To investigate the regulatory function of these two eCREs, we first performed chromosome conformation capture combined with high- throughput sequencing (4C-seq) that anchored at rs79423518-tagged CRE, and observed a strong interaction between the CRE and the promoter region of *PRDX5* and *TRMT112*. This suggests that a direct regulation between rs79423518-tagged CRE and the promoter of two target genes. Then, luciferase reporter assay revealed that rs4930698-tagged CRE exhibited both promoter and enhancer activities, and showed an allele-specific effect, while rs79423518-tagged CRE was also found to have regulatory functions (Fig. 2E). By contrast, more pLD signals (29.8%) received multiple causal eCREs that regulated different target genes (Fig. 2B), highlighting the complexity of genetic regulation in highly linked loci. In summary, our endogenous perturbation screen in pLD serves as a valuable method to facilitate the identification of true causal variants and their associated CREs in cases where statistical fine-mapping faces challenges, while also nominating potential target genes regulated by functional CREs. Consistent with recent exogenous research (*10*), our findings question the assumption that a single variant typically accounts for the causality of an independent association locus. Overall, this result underscores the importance of recognizing the complexity of genetic regulation when interpreting GWAS signals.

### Comparison of regulatory effects between eCRE perturbation and conventional eQTL mapping

To assess the physiological relevance of the eCRE-perturbGene pairs uncovered through our multiplexed single-cell CRISPR perturbations, we compared them with the whole blood eQTL mapping results from GTEx V8 and eQTLgen (*28*). Among all significant eCRE-perturbGene pairs identified in pLD regions, approximately 30% associated eQTL-eGene pairs were found to exist in either GTEx or eQTLgen (Fig. 3A). Similar trends were also observed for perturbGenes (Fig. 3B), suggesting that a large proportion of significant hits from CRISPR perturbations were not captured by current eQTL mapping in whole blood. To explore this further, we compared our findings to a larger eQTL dataset from QTLbase (*3*), which integrates multiple eQTL studies from various tissues/cell types and conditions. We found that a large proportion of additional overlaps could be recovered (Fig. S5A-S5D), suggesting different types of CRISPR perturbation could capture some eCREs that are specific to the cell type or environmental condition not being well studied.

**Fig. 3.**
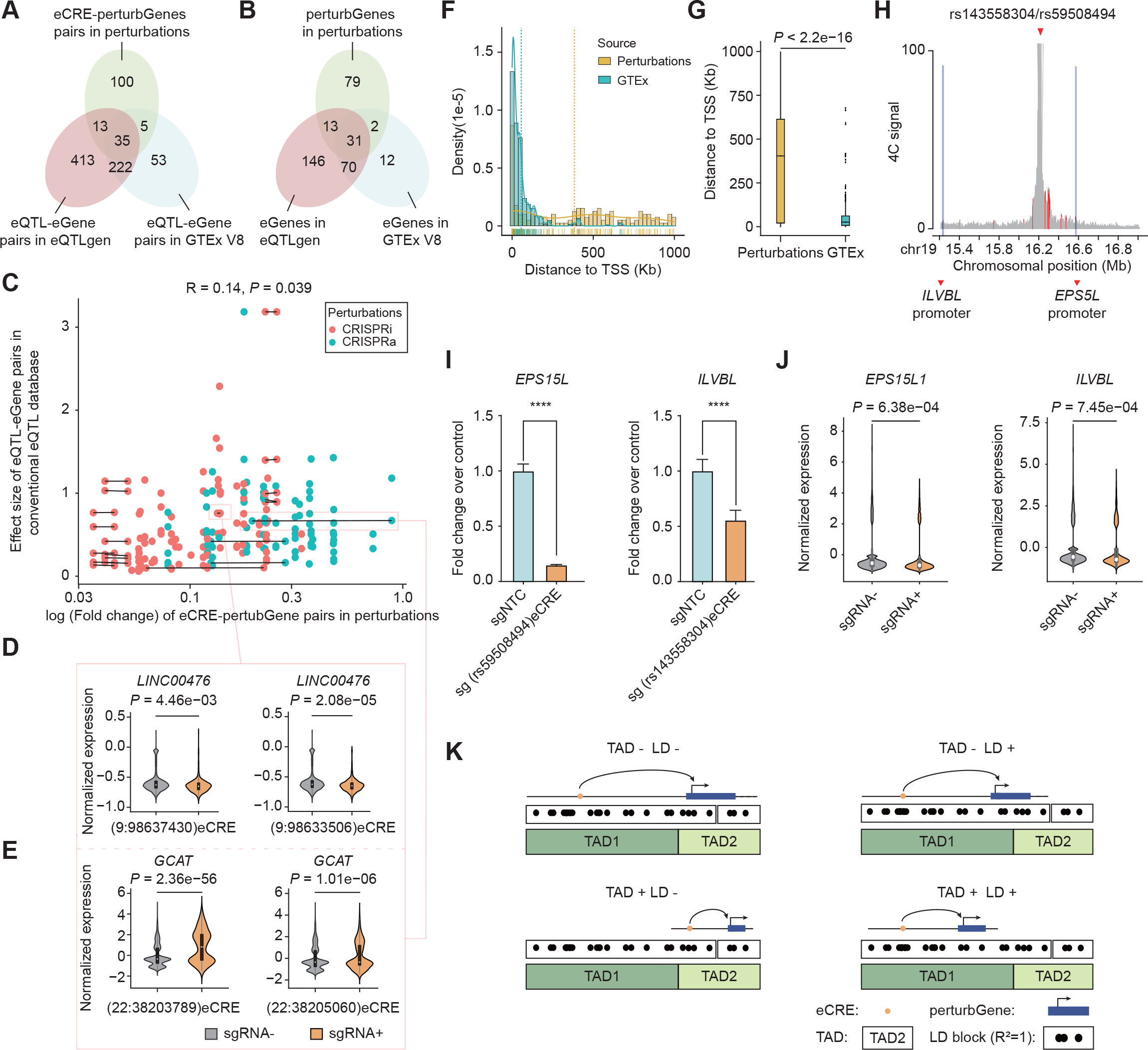
**Illustration and comparison of regulatory effects between eCRE perturbations and eQTL mapping. a**, Venn diagram comparing eCRE-perturbGene pairs identified by the multiplexed single-cell perturbations with eQTL-eGene pairs identified in GTEx V8 and eQTLgen datasets. **b**, Gene-level comparison including perturbGene and eGene, similar with **a**. **c**, Consistency of the estimated effects between eCRE-perturbGene pairs identified by the multiplexed perturbations and eQTL-eGene pairs derived from the conventional eQTL datasets. The red and green dots represent eCRE-perturbGene pairs in CRISPRa and CRISPRi perturbations, respectively, and the lines between the dots indicate that the eCREs are located within the same pLD signal. **d** and **e**, Examples of the varied regulatory effects on the target gene expression for the causal eCREs in two LD signals. **f**, Distribution of the distance between the LEVs and TSS of target genes in multiplexed perturbations (yellow) and GTEx V8 dataset (blue), with the median indicated by dashed lines. **g**, Comparison of the distance distribution between multiplexed perturbation results and GTEx V8 dataset in **f**, with statistical analysis by two-tailed t-test. **h**, 4C results showing the chromatin interaction frequency between the fragment containing rs143558304 and rs59508494 (top red arrow, 4C viewpoint) and the promoter regions (blue shaded region) of the two perturbGenes (bottom red arrows). **i**, RT-qPCR validation results for the sgRNAs targeting eCREs tagged by rs59508494 and rs143558304, with sgNTC as control. Error bars represent standard deviation of the mean. \*\*\*\**P* < 0.0001. **j**, Differential perturbGene expressions in the multiplexed CRISPR perturbations for the causal eCREs tagged by rs59508494 and rs143558304. **k**, Schematic of patterns for causal eCREs regulating their target genes when considering the ranges of TAD and LD block. The symbols representing TAD, LD block, eCRE and pertubGene are shown in the figure.

Previous statistical methods for causal eQTL fine-mapping mostly assumed that functional regulatory variants are sparsely distributed and the tight linkage among them has limited the ability to accurately estimate the magnitude of genetic effects (*4*). Despite the positive overall effect correlation between eCRE-perturbGene pairs from CRISPR perturbations and corresponding eQTL-eGene pairs from GTEx whole blood tissue (R = 0.14, *P* = 0.039), we observed many discrepancies at same pLD signal with multiple causal eCREs (Fig. 3C). For examples, independent CRISPR perturbations showed similar or varied effects for two causal eCREs in pLD respectively (Fig. 3D and Fig. 3E). However, at these multisite regulation pLD regions, the true effect sizes of individual causal loci were indistinguishable (e.g., overestimation or underestimation) using conventional eQTL mapping. These suggest that our multiplexed single-cell CRISPR perturbations offer a more comprehensive assessment of the magnitude of expression-modulating causal effects at endogenous genomic environment.

The high enrichment of functional eQTLs near the transcriptional start site (TSS) had been extensively documented (*29, 30*). By evaluating the target gene TSS distances of eQTL-associated LEVs from GTEx and causal eCRE-associated LEVs from CRISPR perturbation, we found that the LEVs from CRISPR screen hits lie at greater distances from the nearest TSS (median 26 Kb) compared to LEVs in conventional eQTL mapping (median 400 Kb) (*P* < 2.2e-16, T test, Fig. 3F and Fig. 3G, see details in Methods). For instance, rs59508494 (19:16211630-A-G) and rs143558304 (19:16213697-T-TA) are located close to each other (∼2 Kb) within pLD region. The two LEVs are significantly associated with *TPM4* gene expression in GTEx whole blood tissue. Interestingly, Both CRISPRi perturbation screen and RT-qPCR revealed that rs59508494 could regulate a distal gene *EPS15L* (390 Kb) and rs143558304 could regulate another distal gene *ILVBL* (900 Kb), respectively (Fig. 3I and 3J). The long-distance interactions between the two LEVs and their previously unknown targets were validated by 4C-seq that anchored at a genomic fragment containing the two variants (Fig. 3H). Given the evidence that GWAS hits are further from TSSs than eQTLs and show limited overlaps with them (*5–7*), the multiplexed CRISPR perturbations would provide a comprehensive supplement to study the shared genetic effect between disease/trait-causal variants and functional regulatory sites.

Topologically associating domains (TAD) and LD are two measurements of chromosomal interaction and genome genetic structure respectively, by which the genome is divided into different segments. Boundaries and ranges of both measurements are important for exploring the relationship between genetic architecture and gene regulation. Previous findings revealed that genomic architectures of genetic and physical interactions are generally independent, and the regulation range of eQTL-eGene is irrelevant with LD (*31*). Here, using the eCRE-perturbGenes identified by CRISPR perturbation, we reassessed such relevance and found that the causal eCREs regulating their target genes within the same TAD are more likely to locate in a highly linked LD block (Odds ratio = 14.2, *P* = 9.4e-11, Fisher’s exact test, Fig. 3K, see details in Methods). This also suggests that artificial genetic perturbation would capture additional layers of gene regulation against the traditional eQTL mapping in homeostatic conditions.

### Endogenous perturbation effects are poorly predicted via computational predictions and functional annotations

Given statistical fine-mapping faced the tremendous challenge to accurately identify true causal eQTL variants in pLD, we next sought to evaluate the performance of the existing computational methods and functional annotations in distinguishing the regulatory potential of eCREs through endogenous CRISPR perturbation. First, we leveraged 20 functional/pathogenic variant scores to test their abilities of causal eCRE/LEV classification (Table S6 and Table S7, see details in Methods). Consistent with the previous benchmarks using massively parallel reporter assays (MPRAs) data (*10, 32*), existing prediction tools showed restricted performance in discriminating significant causal eCREs or corresponding LEVs from non-significant ones in our CRISPR perturbation screens (Fig. 4A, Fig. 4B, Fig. S6A and Fig. S6B). Specifically, the results showed that among the 20 methods for predicting functional eCRE/LEV, DVAR (*33*), RegBase_REG (*34*) and Eigen-PC (*35*) achieved a better performance (Area Under the ROC Curves (AUCs) are close to 0.6) than others in the majority of benchmarks. Notably, these top-performed tools were either learned from unsupervised algorithms (such as DVAR and Eigen-PC) or presented an ensemble score by integrating existing prediction methods (like RegBase), implying they could capture unknown features that explain the endogenous activity of regulatory sites. Second, we applied a recent deep learning model, Enformer (*36*), which predicts sequence effects on gene expressions and chromatin states, to investigate the agreement between predicted effects and perturbation effects at tested eCRE/LEV sites (Table S6 and Table S7, see details in Methods). We noted similar enrichment patterns of investigated eCREs and the associated LEVs on Enformer scores, in which significantly casual eCREs or tagging LEVs enriched at the top percentiles of Enformer scores from the first two components, compared with non-significant ones (Fig. 4C, Fig. 4D, Fig. S6C and Fig. S6D). Nevertheless, the differences between the eCRE groups remain small.

**Fig. 4.**
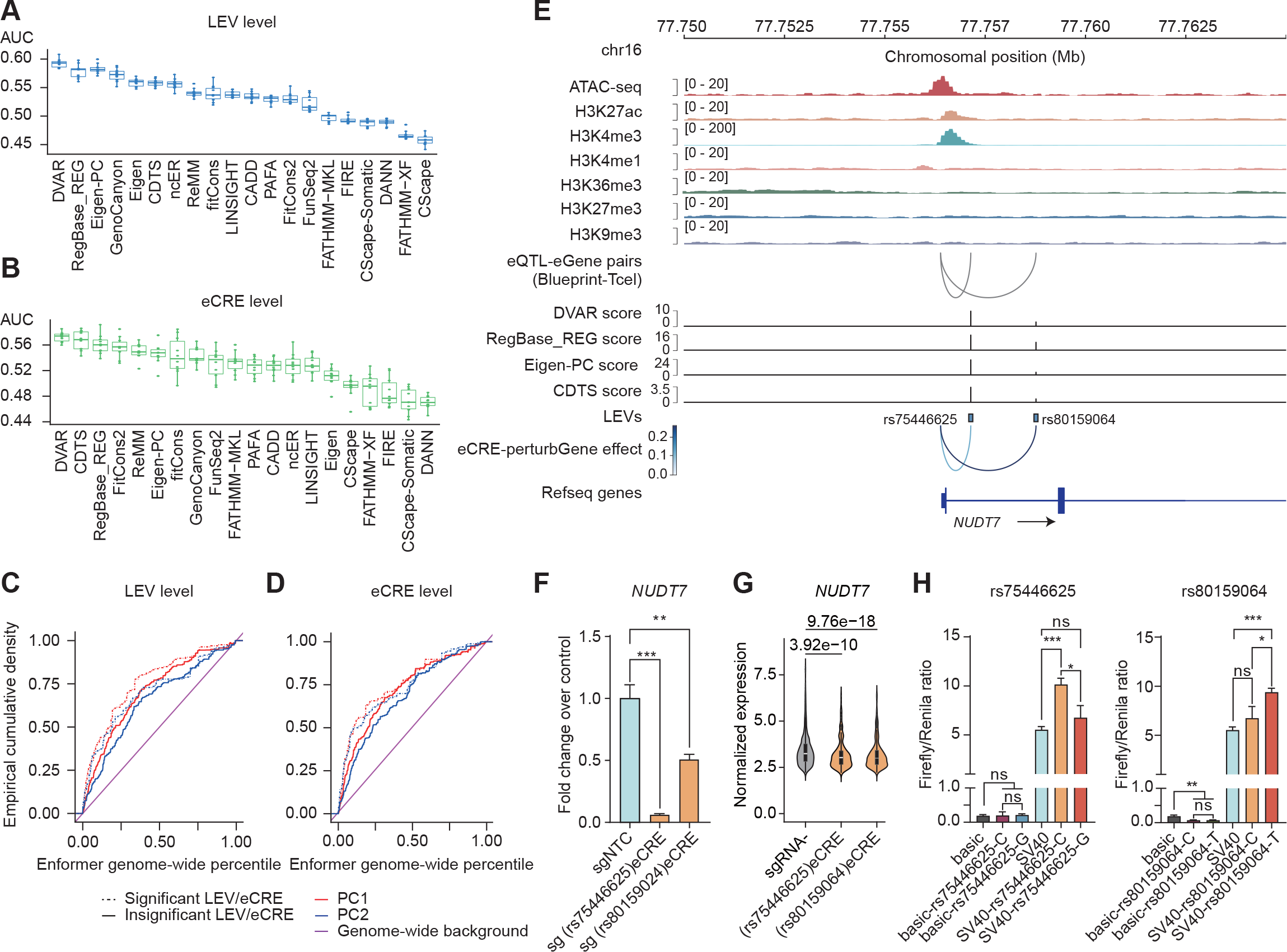
**Evaluation of the endogenous perturbation effects via computational predictions and functional annotations. a**, Benchmarks of existing computational methods in predicting causal eCRE-associated LEVs, non-significant eCRE-associated LEVs were used as negative samples. **b**, Benchmarks of existing computational methods in predicting causal eCRE by taking median prediction scores of all possible ± 1 Kb variants surrounding corresponding LEVs, non-significant eCREs were used as negative samples. **c**, Empirical cumulative probability distribution of the first and second Enformer principal component scores for significant and non-significant eCRE-associated LEVs. **d**, Empirical cumulative probability distribution of the first and second median Enformer principal component scores for significant and non-significant eCRE, by taking median prediction scores of all possible ± 1 Kb variants surrounding corresponding LEVs. **e**, Epigenomic and computational evidence for two causal eCRE-associated LEVs, rs75446625 and rs80159064. **f**, RT-qPCR validation results for the sgRNAs targeting eCREs tagged by rs75446625 and rs80159064, with sgNTC as control. **g**, Differential perturbGene expressions in the multiplexed CRISPR perturbations for the causal eCREs tagged by rs75446625 and rs80159064. **h**, Luciferase assay results of rs75446625/rs80159064-tagged CREs and allele-specific effects of them. \**P* < 0.05; ** *P* < 0.01; \*\*\**P* < 0.001; \*\*\*\**P* < 0.0001; ns, no significant.

Furthermore, we illustrated a case wherein computational predictions and functional annotations poorly worked. Specifically, we observed that rs75446625 and rs80159064 were two perfectly linked variants located in the intronic region of *NUDT7*. While the rs75446625-tagged CRE harbored several active chromatin states (including open chromatin, H3K27ac and H3K4me3), the rs80159064-tagged CRE was completely depleted from classical markers. As expected, rs75446625 was highly scored by four top-performed scores (including DVAR, CDTS (*37*), RegBase_REG and Eigen-PC) and two Enformer components. In contrast, rs80159064 showed very low scores for most of these tools (Fig. 4E). However, our CRISPRi perturbation screens identified that the CREs tagged by the two LEVs can modulate the expression of a common target gene *NUDT7*, which was also successfully validated through RT-qPCR (Fig. 4F). Luciferase reporter assays revealed that the two eCREs both had regulatory functions as an enhancer and showed allelic-specific effects (Fig. 4G and Fig. 4H). Particularly, compared to rs75446625-tagged CRE, rs80159064-tagged CRE exhibited larger effect in CIRSPR screen (Fig. 4E) and equivalent effects at *in vitro* reporter assays, respectively. Taken together, current computational methods for statistical fine-mapping and functional prediction are less actionable for the identification of true causal regulatory variants in high LD.

### Unbiased endogenous perturbation reveals many unmarked regulatory elements

The majority of previous Perturb-seq studies targeting regulatory DNA sequences typically rely on prior knowledge of specific types of CREs, such as enhancers, open chromatin regions, and transcription factor binding sites (*38*). However, in our CRISPRi/a perturbation screen, we did not use any chromatin marks or sequence features to select LEVs and associated eCREs. This approach provided a unique opportunity to unbiasedly learn the regulatory potential of genomic sequences. By integrating seven well-characterized epigenetic marks in HAP1 cells, we were able to classify the identified eCREs into two major groups: marked CREs and unmarked CREs (UREs) (see details in Methods). Active but not repressive chromatin signals, such as open chromatin (ATAC-seq), enhancer/promoter (H3K27ac, H3K4me1, and H3K4me3), and actively transcribed genomic regions (H3K36me3), were prominent at marked CREs. Surprisingly, we found that over 40% of significant eCREs and their associated LEVs lacked classical epigenetic marks almost entirely (Fig. 5A). This suggests the pervasive existence of UREs across the whole genome that may be driven by specific biological conditions.

**Fig. 5.**
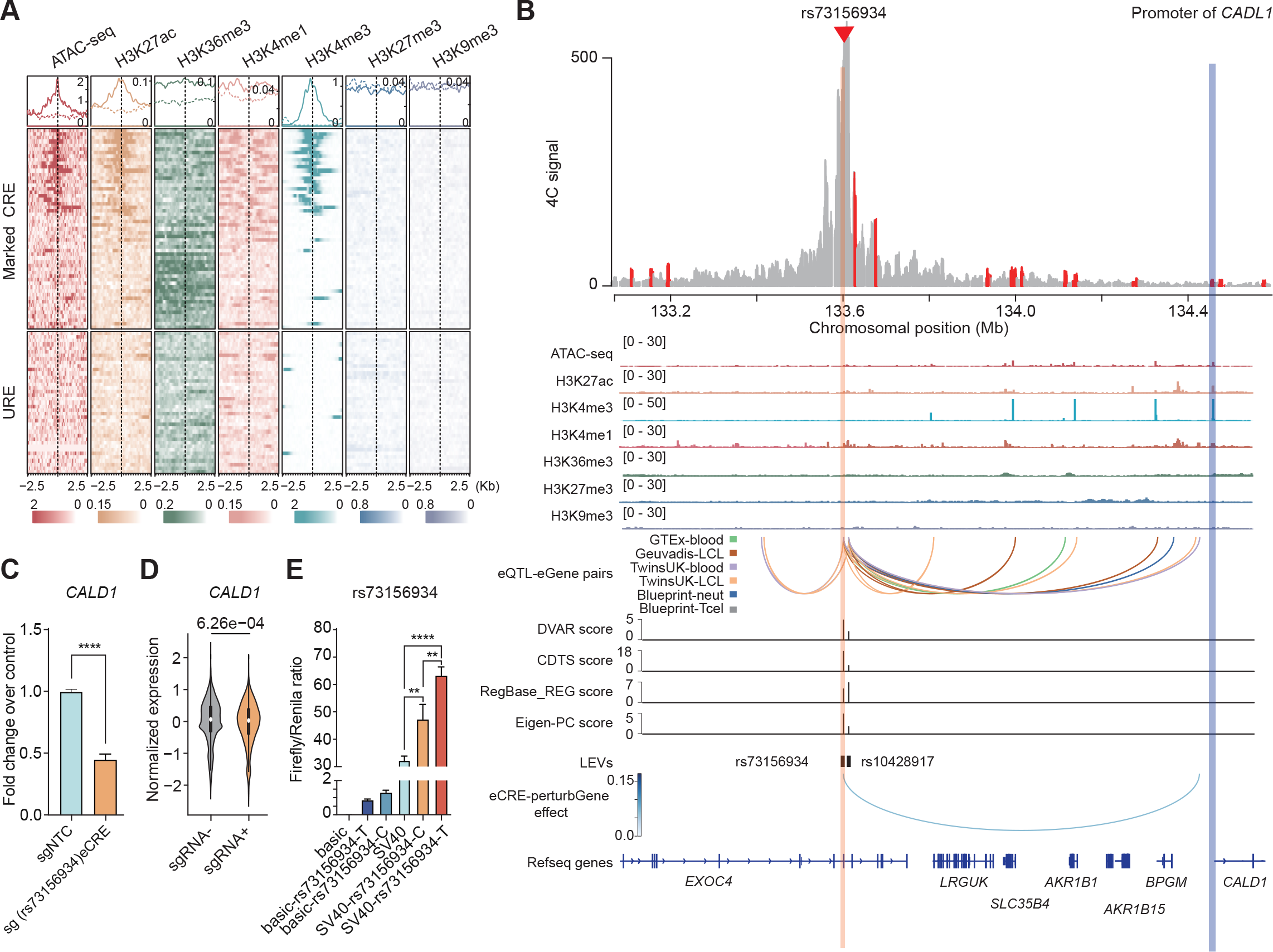
**Evaluation of unmarked regulatory elements revealed by endogenous perturbations and epigenomic marks. a**, The epigenomic profile of HAP1 cells for seven classical marks, including ATAC-seq, H3K27ac, H3K36me3, H3K4me1, H3K4me3, H3K27me3 and H3K9me3, within a ± 2.5 Kb range surrounding causal eCRE-associated LEVs. **b**, 4C results of the interaction frequency between the causal eCRE tagged by rs73156934 (viewpoint, red triangle and yellow shaded region) and the promoter region of *CALD1* (blue shaded region). Epigenomic and computational evidence for two causal eCRE-associated LEVs, including rs73156934 and rs10428917, are shown below. **c**, RT- qPCR validation results for the sgRNAs targeting eCREs tagged by rs73156934 and rs10428917, with sgNTC as control. **d**, Differential perturbGene expressions in the multiplexed CRISPR perturbations for the causal eCREs tagged by rs73156934 and rs10428917. **e**, Luciferase assay results of rs73156934/rs10428917-tagged eCREs and allele-specific effects of them. \**P* < 0.05; \*\**P* < 0.01; \*\*\**P* < 0.001; \*\*\*\**P* < 0.0001; ns, no significant.

To demonstrate the regulatory potential of URE in gene regulation, we performed several functional assays on rs73156934-tagged CRE which showed a significant effect in CRISPRi screen. rs73156934 is an intronic variant at *EXOC4* gene, and its surrounding genomic region is not marked by any epigenetic signals in HAP1 (Fig. 5B) or rarely occupied with H3K36me3 in other tissues/cell types (query from VannoPortal (*39*)), suggesting the regulatory activity of the rs73156934-tagged CRE is preserved in most cellular contexts. Compared to the candidate CREs tagged by other highly linked LEVs (e.g. rs10428917) in pLD region, our CRISPRi perturbation screen and RT-qPCR revealed that this URE could be manipulated to regulate a distal gene *CALD1* (800 Kb) instead of its associated eGene *EXOC4* in GTEx (Fig. 5C and Fig. 5D). The long- distance interactions between the eCRE and *CALD1* were further confirmed by 4C-seq anchored at rs73156934-containing region (Fig. 5B). In addition, luciferase reporter assays indicated that the eCRE had significant regulatory functions and was affected by different alleles of rs73156934 (Fig. 5E). Moreover, the regulatory relationship between rs73156934 and its target gene *CALD1* was supported by several blood single-cell eQTL studies (Fig. 5B).

### Combinatory analysis of CRISPRi/a effects suggests regulatory plasticity of the human genome

Applying both CRISPRi and CRISPRa to the same sgRNA library in the multiplexed single-cell screens enabled us to systematically compare the regulatory effects by different types of perturbations. We found that a large number of perturbations unexpectedly affected their target gene expression in proximity (± 1 Mb surrounding TSS), and the casual eCREs received varied regulatory outcomes under the same perturbation (Fig. 6A). Specifically, CRISPRi can up-regulate nearby target gene expression among one-third of the significant eCRE-perturbGene pairs, and CRISPRa also could down- regulate local gene expression occasionally, although we cannot figure out which hits are from *trans*-effects of perturbation. Besides, over 35% of casual eCRE showed opposite effects on different target genes through either CRISPRi or CRISPRa perturbations. Interestingly, we also revealed 13 eCRE-perturbGene pairs were significant in both CRISPRi and CRISPRa screens (Fig. 6B).

**Fig. 6.**
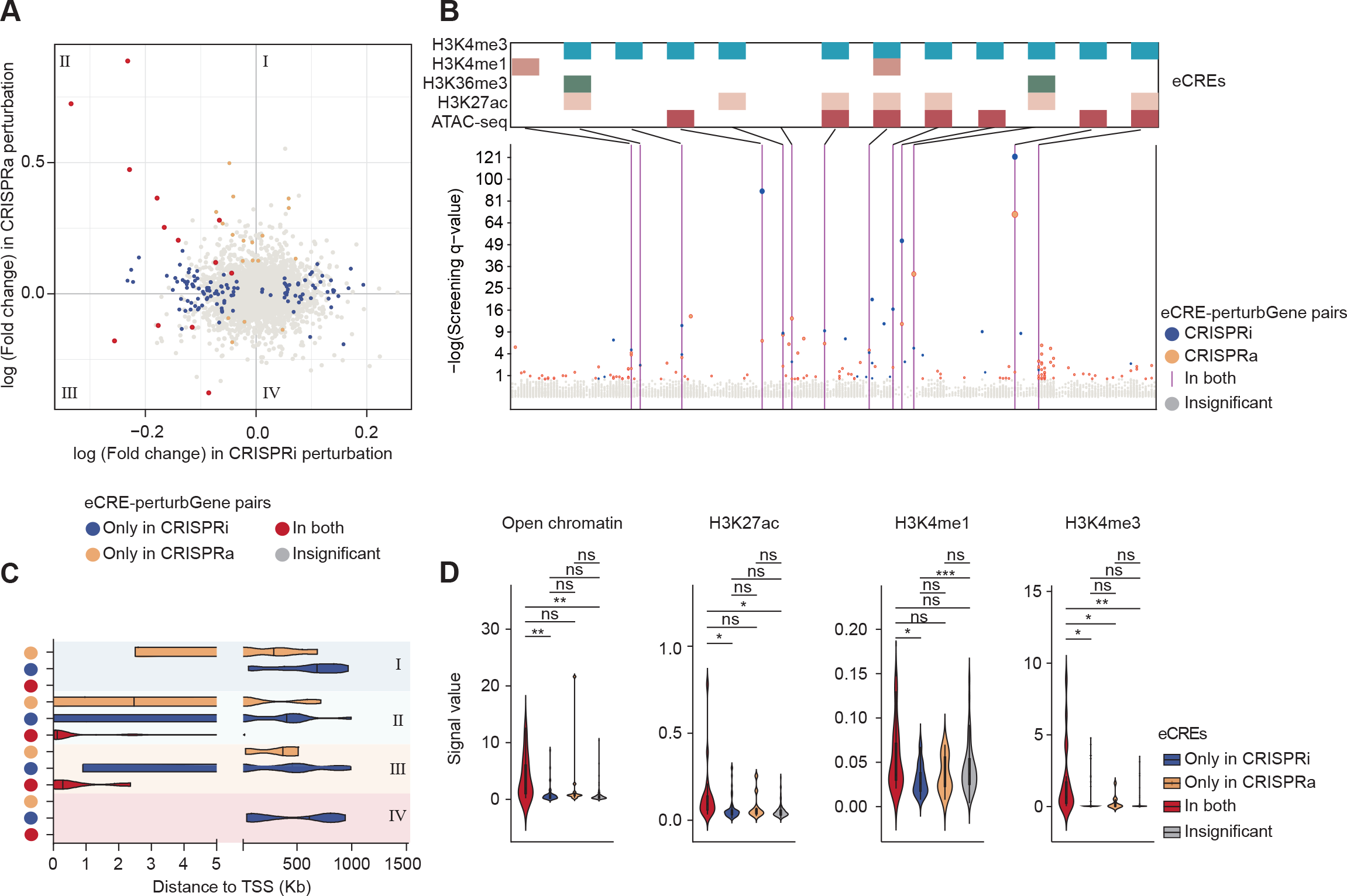
**Genome-wide comparison of various CRISPRi and CRISPRa effects. a**, Magnitude and direction of perturbation effects for all tested eCRE-gene pairs in CRISPRi and CRISPRa perturbations, with blue dots indicating significance only in CRISPRi, yellow dots indicating significance only in CRISPRa, red dots indicating significance in both CRISPRa and CRISPRi, gray dots indicating no significance in both perturbations. Significance threshold is q < 0.2. **b**, LocusZoom-like plot for all tested hits and epigenomic marks for the causal eCREs identified via both CRISPRi and CRISPRa perturbations, with blue indicating significance in CRISPRi, yellow indicating significance in CRISPRa, and the vertical purple line indicating causal eCRE-perturbGene pairs identified by both CRISPRa and CRISPRi perturbations. **c**, Distribution of the distances between causal eCRE-associated LEVs and TSS of perturbGenes in hits at four quadrants obtained from **a**. Left y-axis represents the type of quadrant, with colors consistent in **a**. **d**, Enrichment of epigenomic marks in the four perturbation effect groups, tested by Fisher’s exact test.

By partition all significant eCRE-perturbGene pairs into four categories according to the effect direction of perturbGenes in CRISPRi/a screens (Fig. 6A), we observed that, for the pairs whose gene expression were unexpectedly up-regulated via CRISPRi perturbations (group I and group IV), the distances of casual eCRE from target gene TSS displayed a polarized trend, and most of which were more than 100 Kb away from the TSSs (Fig. 6C). Such phenomenon was also observed for those repressive effects from CRISPRa perturbations (group III) (Fig. 6C). While some effects could be attributed to *trans*-gene regulation, these observations highlight the existence of several hidden but unique mechanisms underlying the distal gene regulation in 3D genome (Fig. 6C). However, the casual eCREs were almost located at the promoter region of their perturbGenes for the 13 significant eCRE-perturbGene pairs identified by both CRISPRi and CRISPRa (Fig. 6C). Besides, compared with casual eCREs detected in single type of perturbation or insignificant ones, more active chromatin signals (including H3K27ac, H3K4me1, H3K4me3 and open chromatin) were enriched at casual eCREs identified by both perturbations. (Fig. 6D). Together, these results indicate that the human genome and chromatin display high plasticity in response to various stimulations at different genomic positions, warranting further research to elucidate the role of sequence variations and chromatin dynamics in shaping functional molecular phenotypes.

## Discussion

Investigating the molecular phenotypes influenced by genetic loci that subsequently lead to the development of complex traits and diseases is a crucial scientific question in the current field of genetics research (*3*). Gene expression, as an essential molecular process for transmitting genetic effects, has made the integrated analysis of eQTL a standard approach to exploring the causal genetic mechanisms of complex traits and diseases (*40, 41*). However, challenges persist in identifying the true causal regulatory variants and their target genes in highly linked regions due to factors such as LD contamination and the complexity of CRE-target gene regulation. In this study, we innovatively employed multiplexed single-cell CRISPRi/a perturbations to investigate the regulatory patterns of genetic loci on target genes in endogenous cellular environments and under pLD conditions. We systematically demonstrated the widespread presence of multiple causal regulatory loci within pLD regions and the intricate nature of their regulatory interactions with target genes. Additionally, our findings revealed that endogenous perturbations can unveil elusive genetic effects and gene expression regulatory patterns not easily detected by traditional eQTL mapping, including evidence for long-range regulatory relationships and high-resolution analysis of regulatory effects. Furthermore, we identified that existing computational methods face difficulties in precisely predicting the influences and consequences of numerous endogenous perturbations on gene expression. We also discovered that approximately half of the casual eCREs lack conventional epigenetic markers, potentially affecting gene expression via distinct regulatory mechanisms. Lastly, through a comparative analysis of CRISPRi/a perturbation effects, we expounded upon the regulatory plasticity of the human genome from a novel perspective. We propose that incorporating multiplexed single-cell CRISPR perturbations into molecular trait QTL and genome-wide GWAS causal variant fine-mapping could complement the limitations of traditional diverse MPRA approaches in assessing the magnitude of genetic effects in endogenous chromatin environments, and their target genes (*8–10*). Our novel findings and supporting evidence will also promote the development of new technologies and theories in functional genomics and related computational biology, ultimately leading to a more comprehensive understanding of gene expression regulatory patterns.

The functional evaluation of the genetic effects for GWAS/eQTL causal regulatory variants by *in situ* modulation of the genomic sequence under endogenous chromatin environments remains challenging. First, unbiased high-throughput screening at single-base resolution remains limited due to the characteristics of editing technologies. For instance, while cytosine base editors (CBEs) and adenine base editors (ABEs) have been widely employed for screening of allele effects under complex phenotypes or at the single-cell level (*11, 12, 42, 43*), their base editing types and uncertain editing outcomes within the editing window hinder their application in functional fine-mapping studies.

Precise editing tools, such as prime editing (PE) and other retron-based systems (*13, 44, 45*), have been used for high-throughput screening of functional single-base variants, but their relatively low editing efficiency in mammalian cells significantly restricts large-scale genome-wide screening. Second, most of the current Perturb-seq-based functional genomic effect assessments are based on diploid cells with heterozygous genetic background (*21, 46*), such as K562 cells. However, in polyploid mammalian cells, incomplete editing or interference of other non-homozygous alleles may mask the expected genetic effects and phenotypes, potentially leading to reduced statistical power. To balance the advantages and disadvantages of existing technologies, we combined CRISPRi and CRISPRa to perform multiplexed single-cell perturbation screening on diploid HAP1 cells, which have a relatively homozygous genetic background.

Previous high-density GWAS, WGS-based eQTL studies, and simulation analyses have indicated that multiple causal effects within specific genomic loci are not uncommon, and multi-target regulation, along with LD contamination, further complicates the fine-mapping of true causal variants and estimation of true effect sizes (*2, 4, 47*). By merely interfering with a large number of potentially independent eQTL regions harboring multiple causal effects using CRISPRi/a, we discovered that 19.3% of these regions contain multiple causal eCREs targeting the same target gene. Consistent with previous MPRA in high LD (17.7% of eQTLs exhibit more than one major allelic effect) (*10*), our results confirm the widespread presence of multisite regulation of gene expression under endogenous genetic systems, emphasizing the serious need to consider such situations when fine-mapping causal variants in different LD regions. Additionally, through CRISPRi/a screening, we found that causal eCRE(s) within the majority of pLD signals (around 80%) can distally impact the expression levels of multiple target genes and are not necessarily always the nearest genes. Compared to traditional eQTL mapping, which more readily detects signals near TSS (*5, 6*), single-cell CRISPR-based QTL mapping approach may be better suited for interpreting missed signals in GWAS-eQTL colocalization studies. Consequently, the systematic integration of large-scale single-cell CRISPR-based eQTLs and traditional context-specific eQTLs will further unravel the ’missing regulation’ phenomenon in non-coding regions of GWAS (*7*).

Fine-mapping computational methods based on functional annotation have been widely used to explore GWAS/QTL causal variants and their potential functions (*48*). However, both our work and several current studies have observed that existing computational methods and bioinformatics tools struggle to accurately predict the functional consequences (e.g., expression effects) of a given DNA sequence or variant allele (*10, 32*), and the reasons for these inconsistencies remain unexplained. The most plausible explanation might be that the non-coding regulatory features used in current prediction models are largely similar, lacking a novel perspective on how regulatory sequences exert their functions; moreover, the functional genomic features used for computational modeling are predominantly measured in normal tissue or cellular environments, with insufficient data under diverse biological conditions. By integrating epigenetic information, we have revealed that approximately half of the significant causal eCRE genomic regions are severely lacking traditional epigenetic markers (referred to as UREs), and some loci even lack epigenetic markers across all known biological conditions. UREs have been considered functional in previous GWAS and functional genomic studies (*9, 15, 16, 49, 50*), which drives researchers to further investigate universal regulatory mechanisms to explain their regulatory potential, and provides a direction for improving the performance of prediction methods.

In various types of perturbations, we found that CRISPRi has better effects than CRISPRa in driving changes in gene expression. Recent research has discovered that the efficacy of CRISPRa depends on basal expression and chromatin state, and bivalent genes are more sensitive to this perturbation (*51*). However, CRISPRi may be more capable of modifying and altering chromatin structure, inducing heterochromatin formation in any genomic region where transcription occurs, thereby causing changes in the expression of multiple genes in close chromosomal proximity. Therefore, introducing a combination of dCas9 fused to different activation domains, such as enCRISPRa technology (*50*), may enhance the power of functional eCRE detection. On the other hand, we found that a portion of CRISPRi/a perturbations exhibited opposite trends to the expected gene expression disruption effects. Presumably, the effects of CRISPRi and CRISPRa on gene expression can depend on the specific regulatory elements and factors present within the targeted genomic region, such as the location and orientation of the targeted site relative to regulatory elements, the activity and accessibility of chromatin and epigenetic modifiers, the availability and activity of other transcriptional regulators, as well as the competition among different CREs (*18, 52, 53*). For example, a genomic region containing both an enhancer and a silencer element might exhibit different responses to CRISPRi and CRISPRa depending on which element is targeted. Additionally, some regions may contain a composite enhancer/silencer element that is responsive to both positive and negative regulatory signals, or they may incorporate multiple distinct CREs that are differentially responsive to transcriptional activators and repressors.

Our study has the following limitations and unresolved issues. First, in order to select reproducible independent eQTL signals, we systematically integrated seven blood-derived WGS-based eQTL datasets and used LD data from 1000 Genomes Project European population to screen for associated pLD LEVs. Although most of the used eQTL samples have European ancestry and were derived from blood tissues, factors such as heterogeneity of WGS variant calling, discrepancies in LD structure among subpopulations, and differences in blood cell-specific gene expression levels may lead to incomplete selection of pLD LEVs in each independent eQTL signal. Second, as emphasized before, we still have difficulty in screening the genetic effects of causal variants on specific phenotypes at the single-base level in a high-throughput and unbiased manner. CRISPRi/a-based genomic perturbation cannot accurately assess the genetic effects at the variant and allelic levels, nor can it distinguish between closely located genetic variations (e.g., less than 1kb). Additionally, some potential functional CREs cannot be driven by CRISPRi/a. Therefore, we need to develop efficient single-cell single-base perturbation technologies to accurately measure the true effects of genetic variations on gene expression. Finally, this experiment only evaluates the effects of genetic loci using limited gene expression as the readout due to genomic distance and expression level constraints. However, many functional regulatory variations may affect gene expression levels through moderate or *trans*-effects or directly influence other transcriptional-level molecular phenotypes without changing gene expression conditions. Despite these limitations, our study reveals several unique patterns for the complexity of gene expression regulation.

## Methods

### Plasmids

The dCas9-VP64-blast (Addgene, #61425) and dCas9-KRAB-blast (Addgene, #89567) plasmids were separately used to perform CRISPRi and CRISPRa experiments. The sgRNA oligos were annealed and cloned into the CROP-seq-opti plasmid (Addgene, #106280) after *Bsm*BI (NEB, R0580L) digestion. The sgRNA library was synthesized by Synbio Technologies. The psPAX2 (Addgene, #12260) and pMD2.G (Addgene, #12259) were used to pack lentiviruses. The plasmids were used for dual-luciferase reporter assay, including the pRL-TK Renilla luciferase control vector (Promega, E2241), pGL3- Promoter (Promega, E1761), and pGL3-basic (Promega, E1741).

### Cell lines and cell culture

293FT (ThermoFisher, R70007) cells were cultured in Dulbecco’s Modified Eagle’s Medium (DMEM; ThermoFisher, 11965092) containing 10% FBS. HAP1 (Horizon Discovery) cells were cultured in Iscove’s Modified Eagle’s Medium (IMDM; ThermoFisher, 31980030). Both cells were supplemented with 10% fetal bovine serum in 5% CO_2_ at 37°C. Diploid HAP1 cells were isolated from Hoechst (MCE, Y-15559)-stained HAP1 cells by flow sorting.

### WGS-based blood eQTL fine-mapping

Seven WGS-based blood-derived eQTL datasets were used to identify independent eQTL signals and fine-mapped LEVs based on CaVEMaN method (*54*). Specifically, the fine-mapped eQTL variants for GTEx V8 whole blood eQTLs (670 whole blood samples) were obtained from GTEx portal (*2*), and the fine-mapped eQTL variants for several other cohorts with blood-derived samples, including Geuvadis (445 lymphoblastoid cell samples), and TwinsUK (523 lymphoblastoid cell samples and 246 whole blood samples), were obtained from the original CaVEMaN publication (*54*). Additionally, for BLUEPRINT samples (190 monocyte samples, 196 neutrophil samples and 165 CD4+ T cell samples), we conducted eQTL mapping using FastQTL (*55*) based on the individual WGS genotypes and normalized RNA-seq quantifications. Permutation test was applied to estimate the nominal *P* thresholds required for the conditional analysis, Then, we used CaVEMaN to perform eQTL fine-mapping for each significant eQTL signal, and extracted the best eQTL that was most likely to be causal.

### Nomination of reproducible pLD signals with multiple LEVs

Based on eQTL fine-mapping results from GTEx V8 whole blood samples, we sought to identify independent eQTL signals that are reproducible in at least one additional blood- derived eQTL dataset. Generally, we measured LD using genotypes of European samples from 1000 Genomes Project phase3 (*56*) and searched perfectly correlated LEVs (R^2^ = 1) with each GTEx LEV among fine-mapped eQTL signals of other blood- derived datasets. Ultimately, we integrated all fine-mapped variants in seven eQTL datasets to retain independent signals which contains at least two undistinguishable LEVs in pLD. To ensure that the genes being tested were measurable in 10x scRNA-seq, we only included top 20% expressed genes in HAP1 cells. Then, LEVs located in protein-coding/splicing-altering region of the human genome or in the chr15 diploid region of HAP1 cell line were excluded. Due to the limitations of CRISPRi/a in the scope of genomic targeting, we only retained signals with a distance greater than 1 kb between each pairwise LEVs. Thus, pLD signals that were distant from these highly expressed genes (> 1 Mb) in HAP1 cells were also excluded. All of these analyses, including the following, were based on the human genome assembly GRCh37/hg19.

### sgRNA library design

We used FlashFry 1.9.3 (*57*) to design sgRNAs targeting each eCRE. We excluded sgRNAs whose splicing sites located more than 50 bp away from the eCRE-associated LEVs and with IN_GENOME>= 2. Then we ranked the remaining sgRNAs based on Doench2014OnTarget, Hsu2013, and Doench2016CDFScore, as well as otCount. We selected the top two sgRNAs targeting each eCRE (except for cases where only 1 or no sgRNA met the criteria). To include appropriate controls, we incorporated 39 non-targeting control gRNAs from the Human CRISPR Knockout Pooled Library (GeCKO V2) (*58*) as negative controls and nine sgRNAs targeting promoters of 4 genes (*EZH2* (*59*), *CANX* (*60*), *NEAT1* (*61*), *PARK7* (*20*), among which *EZH2* was targeted by three sgRNAs) and two sgRNAs targeting an enhancer of *NMU* (*60*) from different studies as positive controls. The oligos of the sgRNA library were synthesized and cloned to the CROP-seq-opti vector after *Bsm*BI digestion by Synbio Technologies, according to the GeCKO V2 (*58*). Additionally, sgRNAs targeting the same eCRE or positive control site are referred to as a “sgRNA group”. In the following bioinformatics analysis, all sgRNA groups that target eCRE are referred to as “perturbative sgRNA groups”, whereas all other sgRNA groups are referred to as “control sgRNA groups”.

### Quality control of synthetic sgRNA library

To assess the quality and potential bias of the sgRNA library, the sgRNA sequences were amplified using PCR from either the plasmid library or genomic DNA extracted from HAP1 cells 4 days post-transduction, using qsgRNA-F and qsgRNA-R primers and 2x Phanta Max Master Mix (Vazyme, P515-01). The resulting PCR products were purified using the QIAquick PCR Purification Kit (Qiagen, 28106) and then used to generate a next-generation library using the VAHTS Universal DNA Library Prep Kit for Illumina V3 (Vazyme, N607-01). The library was purified using AMPure XP beads (Beckman Coulter, A63880) and sequenced on an Illumina NovaSeq PE150. The sgRNAs were identified by matching the sequence to “CACCG[sgRNA]GTTT” and compared to the designed sgRNAs to determine the correct rate of the sgRNA plasmid library. The potential bias was evaluated by calculating the ratio of 90^th^ percentile to 10^th^ percentile sgRNAs that had at least one sequencing read.

### Production of lentivirus

293FT cells were seeded 24h prior to lentivirus packaging. The lentivirus was produced by co-transfecting the backbone plasmid with viral packaging plasmid (psPAX2) and viral envelope plasmid (pMD2.G) at a ratio of 4:3:1 into 293FT cells using LipoFiter (Hanbio, HB-LF-1000) according to the manufacturer’s instructions. The cell culture supernatant was collected 48h post-transfection and filtered using a 0.45 μm filter.

### Construction of dCas9-KRAB and dCas9-VP64 stably expressed HAP1 cells

Lentivirus containing dCas9-KRAB-blast and dCas9-VP64-blast were separately used to construct stably-expressing HAP1 cells. Diploid HAP1 cells were seeded into a six-hole plate supplemented with 8 μg/mL polybrene (Beyotime, ST1380) 24h prior to lentivirus infection. The lentivirus was added to the cells and 24 hours post-infection, blasticidin (ThermoFisher, 461120) was added to the culture supernatant to a final concentration of 10 μg/mL. Selection was maintained for 3 days to obtain stably-expressing cell lines. The Anti-CAS9 Antibody (BOSTER, BM5120) and Anti-β-Actin antibody (ABclonal, AC02) were used to verify the expression of the dCas9-KRAB and dCas9-VP64 by Western blot. The stably-expressing cell lines were then plated in a 96-well plate by limiting dilution and cultured for 2 weeks to obtain single clones. The activation and inhibition efficiency of single clones were verified by infection with lentivirus containing sgRNA targeting the TSS of *EZH2*. The two most efficient clones of each cell line were selected for perturbation.

### Infection of lentivirus with different MOIs

The lentivirus supernatant of the sgRNA library was concentrated using the ViraTrap™ Lentivirus Concentration Reagent (Biomiga, BW-V2001-01) and titrated using the Lenti- Pac™ HIV qRT-PCR Titration Kit (GeneCopoeia, LT005). The dCas9-KRAB and dCas9- VP64 stably expressed HAP1 cell lines were seeded into a 24-well plate and transfected with the concentrated sgRNA library at MOI=500 (moderate MOI). To increase the infection efficiency, 8 μg/mL polybrene was added to the cell culture. After 24 hours, the cells were treated with 0.3 μg/mL puromycin (Sigma-Aldrich, P7255) for three days. For high MOI infections, a second round of infection was performed to achieve a higher MOI.

### scRNA-seq and sgRNA-transcript enrichment

To prepare multiplexed CROP-seq libraries, adherent cells were digested with 0.25% trypsin and collected in a 15mL tube containing serum-containing medium, followed by centrifugation and washing of the cell pellet with serum-free basal medium to obtain a single-cell suspension. The cell density was adjusted with a cell counter and the 10x Genomics Chromium Single Cell 3’ Library reagents V3 were used according the manufacturer’s instructions. To enrich for sgRNA-transcripts, PCR was performed on 15 ng of cDNA from the 3’ single-cell RNA libraries using SI-PCR primer and 10x-sgRNA i7- N720 primer in each 50 μL reaction with an annealing temperature of 60°C and 2 × Phanta Max Master Mix. The enriched sgRNA libraries were sequenced on Illumina NovaSeq6000 PE150 with the same configuration as the standard 10x libraries.

### scRNA-seq data processing

The sequencing data from 10x Genomics Chromium 3’ scRNA-seq underwent initial processing with Cell Ranger v5.0.1, which involved sequence alignment, filtering, barcode counting, and UMI counting. The resulting data were further analyzed using Seurat v4.0.3 (*62*), where a series of quality control measures were applied. Specifically, cells with mitochondrial percentage exceeding 10% or having less than 200 gene UMIs were removed. Additionally, genes that were expressed in less than 0.525% of cells were filtered out. Droplets were also identified and removed using scDblFinder (*63*).

### sgRNA and single cells assignment

With the fore-mentioned amplification protocol (*25*), we enriched sgRNAs and then calculated the distribution of sgRNAs in different perturbation. Firstly, we aligned the enrichment reads to reference genome (GRCh37/hg19) using Cell Ranger v5.0.1, following the same procedure as for scRNA-seq data processing. We then extracted the reads that failed to align to the reference genome using Samtools (*64*). Next, we mapped each unmapped read to the sequences of the sgRNA library and its reverse complement. To limit the sequence before and after the sgRNA sequence, we used ACCG as the left anchor (which is at the end of the U6 promoter) and GTTT as the right anchor. During the sgRNA detection process, we also extracted the cell barcode and sgRNA UMI count for each perturbed cell. The number of sgRNAs per cell and the number of cells bearing each perturbation was calculated based on data from multiplexed single-cell CRISPRi and CRISPRa perturbations with different MOIs.

### eCRE-gene association analysis

For each targeted eCRE, we divided cells into two groups based on whether they contain specific sgRNAs targeting observed eCRE, and evaluated the differential expression of candidate genes within 1 Mb of the eCRE between perturbative sgRNA group and control sgRNA group using Normalisr (*26*). Normalisr is a unified normalization-association framework for statistical inference of gene regulation. It uses an ingenious normalization strategy followed by a regular linear regression model. The normalization step estimates the pre-measurement mRNA frequencies from the scRNA-seq UMI counts and regresses out the nonlinear effect of library size on expression variance. And then a linear regression model tests the associations between eCRE and perturbGene, where two- sided *P* were computed from Beta distribution, and log fold change was estimated using maximum likelihood. We also included additional covariates including mitochondrial percentage, unique gene count, and sgRNA count in the association testing to improve the accuracy of the results. To avoid false positive results, we estimated the *P* of the associations between genes and NTC sgRNAs and considered them background *P*. An empirical *P* was calculated based on the background *P* and raw *P*, which was then adjusted by FDR (q-value) for multiple testing corrections. Finally, we defined a 0.2 threshold of q-value based on the NTC tests as they are subject to the same sources of error as the eCRE-targeting sgRNAs.

### Simulation for power estimation

Based on Splatter (*65*), we simulated synthetic scRNA-seq datasets with various group.prob parameters to mimic CROP-seq cells under different perturbation conditions. The Splatter simulation process consisted of two steps. First, we estimated the necessary parameters for simulation using splatEstimate from the CROP-seq dataset. Subsequently, we utilized the estimated parameters to simulate synthetic scRNA-seq datasets with splatSimulate, where the average expression of each gene was randomly sampled from a gamma distribution and the cell’s experimentally measured count was sampled from a Poisson distribution. To simulate perturbation under different MOIs, we used the *group.prob* parameter, which is a vector containing two values. The first value was obtained by dividing the number of perturbations per cell (ranging from 200 to 3,800) by the total number of cells, and the second value was obtained by subtracting the first value from 1. We used *nCells* parameter to simulate perturbation under different cells (ranging from 5,000 to 20,000). Finally, we assessed the power of each simulation by computing the ratio of recovered CROP-seq results to all CROP-seq results.

### Validation of individual hits

For validation of positive eCRE-perturbGene pairs identified by multiplexed CRISPRi/a perturbations, the expression of perturbGenes was measured by RT-qPCR following individual perturbation with sgRNAs targeting the eCREs. Briefly, oligonucleotides of sgRNAs targeting causal eCREs and random sgNTC were synthesized and annealed, and then cloned into CROP-seq-opti plasmids that were digested by *Bsm*BI. The resulting plasmids containing sgRNAs targeting the same eCRE were mixed in equal amounts and packaged into lentiviruses. The dCas9-KRAB and dCas9-VP64 stably expressed HAP1 cell lines were infected with the lentiviruses and selected with puromycin. RNA was extracted from cells, and cDNA was generated using HiScript II Q Select RT SuperMix for qPCR (+gDNA wiper) (Vazyme, R223-01). The cDNA was amplified with 2x SYBR Green qPCR Master Mix (Bimake, B21202) using ACTB as a positive control. Data were analyzed using the 2^-ΔΔCt^ method, with sgNTC as the control. The effect on gene expression of sgRNAs targeting eCREs was calculated after normalization.

### Luciferase reporter assay

Genomic sequences containing causal eCREs with different alleles of LEVs were amplified from HAP1 cell genomic DNA using overlapping PCR. The resulting fragments were integrated upstream of the luciferase gene in pGL3-Basic and pGL3-Promoter plasmids, and the concentration of the recombinant plasmids was determined using the Qubit dsDNA HS Assay Kit (ThermoFisher, Q32851). 293FT cells were transfected with 1 µg of recombinant plasmids or blank vectors with 40 ng of pRL-TK Renilla luciferase control vector using Lipofiter. After 24 h, cells were lysed with 200 µL lysis buffer and shaken slowly on ice for 10 min. Relative luminescence signals were measured using the Dual-Luciferase Reporter Assay System (Promega, E1960) and GloMax® 20/20 Luminometer by normalizing firefly luciferase signal with renilla luciferase signal.

### Circularized chromosome conformation capture (4C) assay

We used the 4C-seq method as described previously (*66*) to validate eCREs tagged by specific LEVs. To prepare the 4C-seq samples, approximately 1×10^7^ HAP1 cells was collected to crosslink by formaldehyde for 10min, and quenched with glycine at a final concentration of 125 mM. Cell pellets was washed twice with cold PBS and lysis on ice for 15 min with 5 mL lysis buffer. The nuclei were digested with the first enzyme (*Cvi*QI (NEB, R0639L) or *Nla*III (NEB, R0125S)) and ligated with T4 ligase (NEB, M0202) overnight at 16°C. Proteinase K (Transgene, 20 mg/ml) was added to the sample, which was then placed in a 65°C water bath overnight to reverse the cross-links. After de- crosslinking, DNA was extracted using phenol-chloroform-isoamylalcohol (Invitrogen, 25:24:1) and finally dissolved in 150 μL of 10 mM Tris-HCl (pH 7.5). The sample was digested with the second enzyme (*Eco*RI (NEB, R0101S) or *Dpn*II (NEB, R0543L)) and ligated with T4 ligase (NEB, M0202) overnight at 16°C. DNA precipitation was obtained by ethanol precipitation, and the DNA was dissolved in 150 μL of 10 mM Tris-HCl (pH 7.5). The 4C template was obtained by adding 750 μL of Buffer PB solution (QIAGEN, #28106) to the sample and dividing it into three spin columns for centrifugation and washing. Each spin column was washed with 50 μL of 10 mM Tris–HCl (pH 7.5). The eluates from the three tubes were collected as 4C template. Viewpoint-specific amplification was performed with 8 × 200 ng 4C template using 2 × Phanta Max Master Mix. The next-generation library was constructed using the VAHTS Universal DNA Library Prep Kit for Illumina V3 and purified by AMPure XP beads following the manufacturer’s instructions. The 4C-seq library were sequenced on the Illumina NovaSeq 6000 platform, producing pair-end reads of 150 bp. Processing and visualization of 4C-seq data was done using pipe4C and peakC pipeline as previously described (*66, 67*).

### Measurement of eCRE regulatory range across LD block and TAD

We defined the LD blocks as multiple variants in strong LD across a region. Specifically, we measured LD between variants surrounding each fine-mapped signals (500 Kb) based on genotypes of Europeans from 1000 Genomes Project phase3 (*56*), using the LDproxy module of LDlink (*68*). We then merged all genomic positions for variants in strong LD with the fine-mapped eQTL LEVs (R^2^ >= 0.8) to form the LD blocks. To define a gene-regulating region, we considered the region from the CRE (±1 Kb of LEVs) to 5 Kb upstream and 3 Kb downstream of the gene body of the target gene. We obtained TADs from *in situ* Hi-C of the HAP1 cell line (*69*). By constructing a 2 ×L2 contingency table, we performed a Fisher’s exact test to evaluate the nonrandom association between the number of gene-regulating regions that lie in Hi-C TADs and those that lie in LD blocks.

### Benchmarks with computational prediction scores

We conducted an evaluation on perturbation effect of causal eCREs against various computational methodologies for predicting base-wise regulatory potentials of DNA sequence. Multiplexed single-cell CRISPRi/a perturbation results were used as the golden standard to measure prediction performance. Positive samples were significant casual eCRE-associated LEVs, while negative samples were non-significant ones. To conduct a comprehensive evaluation of the performance of these algorithms, we used the AUC metric of the ROC to distinguish positive and negative samples at both LEV- level and eCRE-level. First, we obtained pre-computed base-wise scores of 20 existingg computational methods from regBase V1.1.1 (*34*). Due to the class imbalance of positive and negative samples, we randomly under sampled negative eCREs to match the number of positive eCREs. We repeated each sampling ten times. Second, Enformer (*36*), a deep neural network to predict gene expression levels given genomic sequence, was employed in our comparisons. Two Enformer principal components (PCs), representing a summary of the most important features that contribute to the prediction of gene expression levels, were calculated and used to generate empirical cumulative probability distributions of the first and second PC scores. By comparing these distributions, we determined if the PCs for significant and non-significant eCRE- associated LEVs differed. For eCRE-level benchmarks, we estimated median, mean, and max scores for ±1 Kb of each eCRE-associated LEVs.

### Epigenomics analysis

The raw sequencing reads of HAP1 ChIP-seq (H3K27ac, H3K27me3, H3K36me3, H3K4me1, H3K4me3, H3K9me3) and ATAC-seq were analyzed by the nf-core pipeline (*70*). Meta-profiles of LEV-centered regions (± 2.5 Kb) were generated from the bigWig files by deepTools (*71*). Heatmaps were generated using EnrichedHeatmap (*72*), and narrow peaks were called using MACS2 (*73*). The causal eCREs that intersected with peak(s) for at least one of fore-mentioned marks were classified as marked eCREs, In contrast, the causal eCREs that do not intersect with peak(s) for any marks are referred to as URE.

### Statistical analyses

Statistical analyses were carried out with GraphPad Prism 8.0 (GraphPad Software). All experiments were performed at least three replicates, unless otherwise noted. Differences in means were compared using an unpaired two-tailed Student’s t-test, and graphed as the means ± standard deviations (SD). Statistical significance denoted as follows: *, *P* < 0.05; **, *P* < 0.01; ***, *P* < 0.001; ****, *P* < 0.0001; ns, no significant.

## Acknowledgements

The work was supported by the following grants: the National Natural Science Foundation of China (32270717 and 32070675 to M.J.L.).

## Author contributions

M.J.L. designed the studies and wrote the manuscript. K.Z., and Y.Z. performed the experiments, analyzed data and wrote the manuscript. C.Y.W., J.H.W., H.C.Y., X.C., L.Z., W.W., X.L.C., and X.F.Y. conducted the bioinformatics analysis or valuation experiment. Y.P.C., M.X.L., W.G.L., K.X.C., and P.C.S. contributed scientific expertise and reviewed the manuscript. All the authors read and approved the manuscript.

## Competing interests

The authors declare no competing interests.

## Data availability

All sequencing data, including scRNA-seq and 4C-seq, generated in this study have been deposited in the Gene Expression Omnibus (GEO) database under accession GSEXXXX. All public data sources and primer sequences used in this study are listed in Supplementary Tables. The average expression of each gene in HAP1 cells in this study was calculated from several bulk RNA-seq data from GEO database under accession number GSE75515, GSE110142, and GSE111272. TAD information of *in situ* Hi-C for HAP1 cells was obtained from GEO under accession number GSE74072. The raw sequencing reads of HAP1 ChIP-seq (H3K27ac, H3K27me3, H3K36me3, H3K4me1, H3K4me3, H3K9me3) were downloaded from ENCODE repositories. Raw data of HAP1 ATAC-seq was downloaded from GEO under accession number GSE111047.

## Notes

### Competing Interest Statement

The authors have declared no competing interest.

